# A large-scale method to measure the absolute stoichiometries of protein Poly-ADP-Ribosylation

**DOI:** 10.1101/2025.03.27.645734

**Authors:** Peng Li, Yajie Zhang, Yonghao Yu

**Affiliations:** Department of Biochemistry, University of Texas Southwestern Medical Center, Dallas, TX 75390, USA; Department of Molecular Pharmacology and Therapeutics, Columbia University Vagelos College of Physicians and Surgeons, New York, NY 10032, USA

## Abstract

Poly-ADP-ribosylation (PARylation) is a reversible posttranslational modification that occurs in higher eukaryotes. While thousands of PARylated substrates have been identified, the specific biological functions of most PARylated proteins remain elusive. PARylation stoichiometry is a critical parameter to assess the potential functions of a PARylated protein. Here, we developed a large-scale strategy to measure the absolute stoichiometries of protein PARylation. By integrating mild cell lysis, boronate enrichment and carefully designed titration experiments, we were able to determine the PARylation stoichiometries for a total of 235 proteins. This approach enables the capture of all PARylation events on various amino acid acceptors. We revealed that PARylation occupancy spans over three orders of magnitude. However, most PARylation events occur at low stoichiometric values (median 0.578%). Notably, we observed that high stoichiometry PARylation (>1%) predominantly targets proteins involved in transcription regulation and chromatin remodeling. Thus, our study provides a systems-scale, quantitative view of PARylation stoichiometries under genotoxic conditions, which serves as invaluable resources for future functional studies of this important protein posttranslational modification.

## Introduction

Mammalian cells are continuously exposed to genotoxic stress and, as a result, have evolved sophisticated mechanisms to detect and signal the presence of damaged DNA, thereby facilitating efficient DNA repair processes. One of the earliest cellular responses following exposure to genotoxic stress occurs through the reversible PTM poly-ADP-ribosylation (PARylation), which is catalyzed by a family of enzymes known as poly(ADP-ribose) (PAR) polymerases (PARPs)^1^. Of all the PARP family members, PARP1 exhibits the highest expression level and has robust enzymatic activity. PARP1 accounts for approximately 90% of PAR polymer formed under genotoxic conditions^2^. PARP1 utilizes intracellular NAD^+^ to synthesize PAR polymers, which are covalently linked to proteins via the attachment to a variety of amino acid acceptors, including Glu, Asp, Ser, Tyr, Cys and Lys^3^.

The synthetic lethality between PARP1 inhibition and *BRCA1/2* deficiency provides the mechanistic foundation for the development of PARP1 inhibitors for the treatment of human malignancies. Indeed, PARP1 inhibitors are successful in the clinic, with four PARP1 inhibitors (olaparib, niraparib, rucaparib and talazoparib) approved by the FDA to treat *BRCA*^mut^ breast, ovarian, prostate and/or pancreatic cancers^4–7^. Despite the tremendous progress of these PARP1 inhibitors, their fundamental mechanisms of action remain incompletely understood.

Auto-PARylated PARP1 functions as a scaffold that facilitates the recruitment of various DNA repair machineries, such as base excision repair, nucleotide excision repair, and double-strand break repair, often through their PAR-binding domains^8–10^. While the initial focus of this field centered on the role of PARP1 in DNA damage repair, our understanding of the functions of PARylation signaling has significantly expanded in recent years. Notably, new findings point to the diverse functions of PARylation signaling in chromatin regulation, transcription, RNA biology, metabolism and viral infections^11, 12^. Furthermore, the recent development of PARylation proteomic technologies has led to the identification of many PARylated proteins, which, in turn, greatly facilitates the functional studies of this important protein modification^13^.

Stoichiometry (the fraction of a given protein that is modified at a given time) is a critically important parameter when assessing the potential function of a posttranslationally modified protein^14^. For example, the transcription factor FOXO3a is known to be phosphorylated by a Ser/Thr kinase Akt, at three critical residues (i.e. T32, S253 and S315)^15^. When Akt is inhibited, FOXO3a is dephosphorylated, and is subsequently translocated into the nuclei to mediate the transcription of a variety of pro-apoptotic genes. It has been shown that the stoichiometry of the phosphorylation at the three aforementioned residues govern the relative distribution of FOXO3a in the cytoplasm vs. nuclei, and in so doing, provides a mechanism for fine-tuning its transcriptional output^15^. Similarly, in the context of PARylation, it has been shown that PARP1 covalently modifies p53 on three evolutionarily conserved residues (i.e., E255, D256 and E268). PARylation of p53 blocks its interaction with the nuclear export receptor, CRM1, resulting in the nuclear accumulation of p53, and initiation of the p53-dependent apoptosis program^16^. It is conceivable that PARylation stoichiometry could be a key factor that regulates the nuclear-cytoplasmic distribution of p53. PARylation stoichiometry is a key parameter when assessing its functional role in regulating protein-protein interactions, subcellular distribution, and protein activity.

While several biochemical methods have been employed to measure stoichiometries of other PTMs (e.g., phosphorylation), these methods are not applicable to PARylation stoichiometry measurement. This is primarily due to the labile, heterogenous and low-abundance nature of PARylation^17, 18^. In particular, PARylation is highly dynamic and is known to be under precise spatiotemporal control^19^. Previously, we developed a large-scale mass spectrometric approach for the site-specific characterization of the Asp-/Glu-PARylated proteome. Coupled with SILAC (Stable isotope labeling by amino acids in cell culture) labeling, this approach allowed the quantification of Asp-/Glu-PARylation events under different cellular states (i.e. control vs. PARP1-inhibited)^13^. However, these relative changes alone cannot be used to determine the absolute PARylation stoichiometries. For instance, a 2-fold downregulation of a PARylation site, as measured in a SILAC experiment, could result from a change in stoichiometry, e.g., from 0.2% to 0.1%, or from 100% to 50%. These two scenarios likely have very different biological implications.

In this study, we developed a large-scale approach for the measurement of absolute PARylation stoichiometries. By combining mild cell lysis conditions, high efficiency boronate enrichment of the PARylated proteins and carefully designed titration experiments, our approach allowed the determination of PARylated stoichiometries of 235 proteins under oxidative DNA damage conditions. Importantly, this approach encompasses all PARylation events occurring on different amino acid side chains, including both known and yet-to-be-discovered acceptors. Our findings reveal a wide range of PARylation occupancy spanning three orders of magnitude (ranging from 35.480% to 0.011%). However, it is noteworthy that the majority of PARylation events occur at low stoichiometries, with a median value of 0.578%. Importantly, our bioinformatic analysis indicates that proteins with high PARylation occupancy (stoichiometry >1%) primarily comprise transcription regulators and chromatin remodelers. This work not only provides valuable insights into the principles of PARylation-dependent signaling, but also seeds the future functional studies of this important protein modification.

## Results

### Overall strategy for determining global PARylation stoichiometries

PARylation is a labile, heterogeneous and low-abundance PTM. We previously developed an integrated strategy towards the site-specific analysis of the cellular Asp-/Glu-PARylated proteome. The bond between ADP-ribose and an Asp/Glu residue constitutes a high-energy “hemiacetal ester”. This moiety is chemically unstable, and is uniquely susceptible to harsh experimental conditions, including heat and high/low pH environments^13^. Furthermore, PARylation also could be aberrantly added/removed by PARP1/PARG during the cell lysis step. In our approach, we used a mild lysis buffer (e.g., a neutral SDS buffer) to deactivate the PARP/PARG enzymes and to universally preserve the PAR chains. Importantly, the SDS buffer system is fully compatible with the subsequent boronate enrichment procedures^13^.

The boronate enrichment strategy leverages the formation of ester bonds between boron and a 1,2-*cis*-diol moiety within ADP-ribose. Here we further evaluated the enrichment efficiency of the boronate system. We found that a single round of boronate enrichment largely abolished the PAR signal in whole cell lysates. Following the second round of boronate pull-down, no detectable PARylation was observed in the supernatant (Fig. 1A), indicating a complete recovery of the PARylated proteins from crude cell lysates by the boronate beads.

**Fig. 1:**
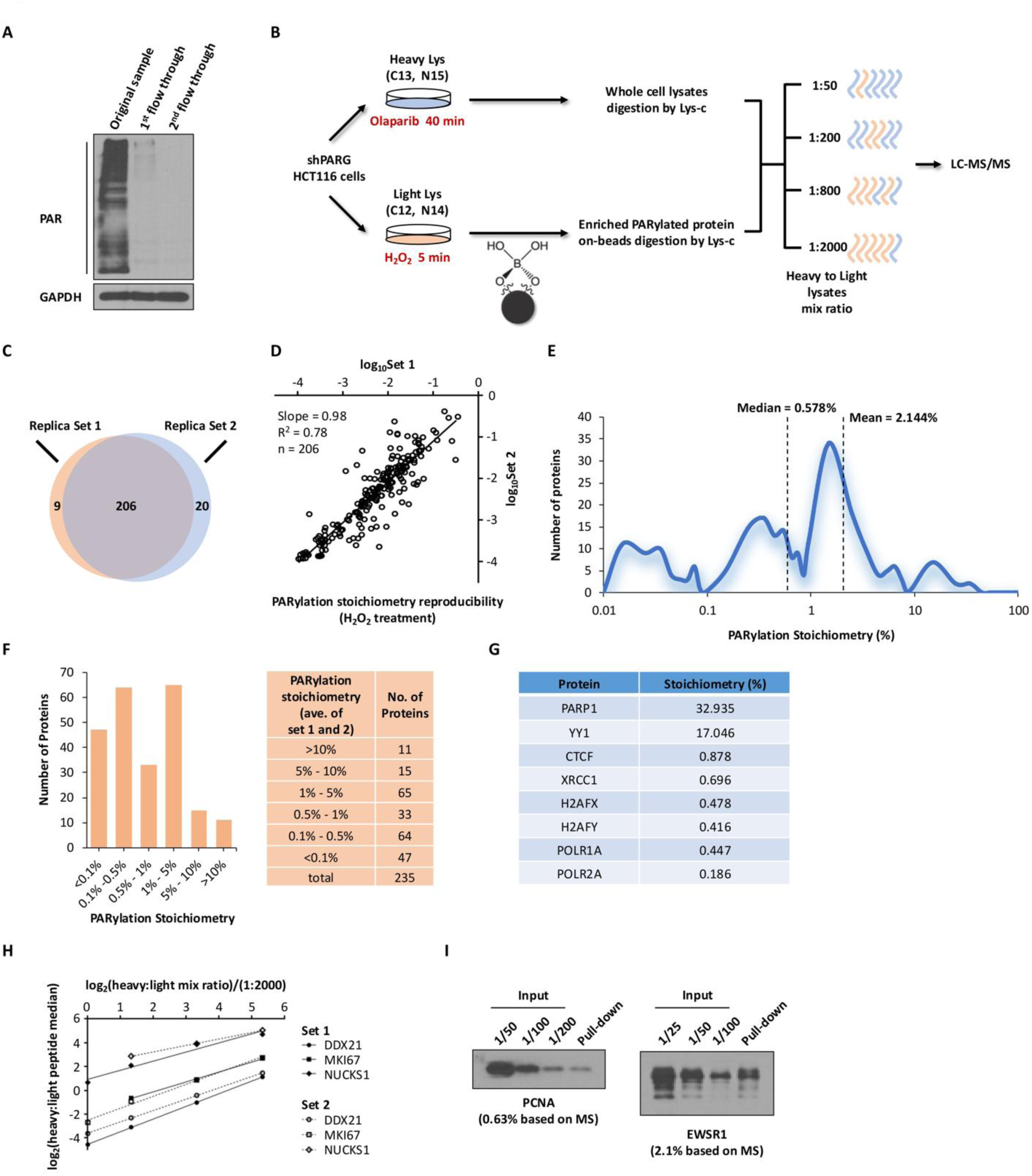
Global PARylation stoichiometry is low but spans large dynamic range. **A** PARylated proteins were quantitatively adsorbed from crude cell lysates using BAB. Cell lysates (shPARG HCT116 cells treated with 2 mM H_2_O_2_) were incubated with BAB twice. The extent of PARylation in the original sample and in two flow-through fractions was assessed using immunoblotting assays with an anti-PAR antibody. **B** Flowchart illustrating the overall strategy for determining global PARylation stoichiometries. Light shPARG HCT116 cells were treated with H_2_O_2_ (2 mM, 5 min) to induce protein PARylation, while heavy cells were treated with Olaparib (1 µM, 40 min) to suppress basal PARylation levels. Enriched PARylated proteins from light cells were subject to on-beads digestion. Light and heavy peptides were then mixed at different protein ratios in the whole cell lysates. **C** Biological replicate experiments were conducted using SILAC HCT116 shPARG cells to determine PARylation stoichiometry. Set 1 identified 215 PARylated proteins, while Set 2 identified 226 PARylated proteins. **D** Reproducibility of PARylation stoichiometry between biological replicates. **E** Overall distribution of protein PARylation stoichiometries (*n* = 235). **F** Distribution of PARylation stoichiometry for grouped PARylated proteins (*n* = 235). **G** PARylation stoichiometry of previously well-studied PARP1 substrates. **H** Dynamic profiles of the median ratio between heavy and light at various SILAC cell lysate mix ratios for individual proteins were compared between biological replicates. **I** Validation of PARylation stoichiometry for PCNA and EWSR1. HCT116 shPARG cells treated with H_2_O_2_ (2 mM, 5 min) were lysed. Lysates were subject to boronate beads pull-down, and the PARylated proteins were eluted as described before. The eluted eluates and the serially diluted cell lysates input were analyzed by immunoblotting to validate PARylation stoichiometry.

We leveraged the excellent enrichment efficiency of the boronate system to develop a strategy towards measuring global PARylation stoichiometries (Fig. 1B). We first generated shPARG HCT116 cells, and label them by stable isotopes (light Lys and heavy Lys, respectively). The light cells were treated with H_2_O_2_, which induced oxidative DNA damage and PARP1 activation^13^. The heavy cells were treated with Olaparib to completely abolish any basal PARylation. We isolated the PARylated proteins from the light cell lysates using the boronate affinity beads^13^. The beads were extensively washed using the SDS buffer, and the PARylated proteins were digested on-beads using Lys-C. On the other hand, the lysates from heavy-labeled cells were directly digested using Lys-C without boronate enrichment. After digestion, aliquots containing light or heavy peptides were combined, and were analyzed by LC-MS/MS experiments. Because the boronate beads completely recover the PARylated proteins from the lysates, we reasoned that the ratio of the light and heavy version of a PARylated protein determined from these SILAC experiments represents its PARylation stoichiometry.

Because PARylation is of very low abundance, it is likely that the PARylated version of a protein presumably represents a very small fraction of the total pool of that protein. The Thermo LTQ-Velos Pro Orbitrap mass spectrometer offers a dynamic range of approximately 5,000^20^. It is important to note that the light/heavy peptide pairs need to have a suitable ratio that is amenable to the dynamic range of the subsequent mass spectrometric detection. Towards this, we performed a series of titration experiments where we used different mixing ratios for the light/heavy samples. For example, we performed on-beads Lys-C digestion of the PARylated proteins isolated from 10 mg of light cell lysates. Separately, we performed Lys-C digestion of 5 µg of heavy cell lysates. The mixing of the peptides resulted in a 1:2000 heavy to light mix ratio. Similarly, a 1:50 heavy to light mix ratio was obtained when the same quantity (10 mg) of light peptides was mixed with the digestion solution of 200 µg of heavy lysates. We performed a series of experiments, employing the different heavy/light mixing ratios ranging from 1:2000 to 1:50 (Fig. 1B). The resulting ratios of the heavy/light peptides were then converted back using the original mixing ratios to determine the PARylation stoichiometry of a PARylated protein.

### Global PARylation stoichiometries span a wide dynamic range

To precisely quantify the absolute stoichiometries of PARylation induced by genotoxic stress, we conducted a large-scale, quantitative MS analysis as described in Fig. 1B. To implement our strategy, we utilized a curated list of PARylated proteins with known PARylation sites from our previous study (Table. S13, Zhang et al., *Nature Methods*)^13^ as a reference. Peptides derived from these proteins were identified, and only the peptides with signal-to-noise ratio (S/N) value > 5 for both heavy and light species were retained.

Next, we calculated the PARylation stoichiometry for each protein based on results obtained from all four mix ratios. Through these experiments, we successfully determined the PARylation stoichiometries of 235 proteins (Fig. 1C). Among them, 215 proteins were identified from Replica Set 1, and 228 proteins were obtained from Replica Set 2 (Supplementary Table 1). A total of 206 proteins overlapped between the two Replica Sets (Fig. 1C). The PARylation occupancy measurements for these 206 overlapping proteins exhibited a robust correlation between replicates (Fig. 1D).

The median and mean PARylation stoichiometries of these PARylated proteins were 0.578% and 2.144%, respectively (Fig. 1E). Merely 11% (26/235) of the proteins exhibited PARylation occupancy greater than 5%, while a remarkable 62% (146/235) of the proteins displayed occupancy levels below 1% (Fig. 1F). These findings suggest a low overall protein PARylation occupancy.

Furthermore, we observed that PARylation stoichiometry varies by three orders of magnitude among these proteins, ranging from 0.011% to 35.48% (Supplementary Table 2). The wide range of PARylation stoichiometries is consistent with PARylation being a highly dynamic protein modification that is regulated in a spatiotemporal manner^21^. This feature again highlights the importance of employing different mix ratios to generate SILAC peptide pairs with suitable dynamic ranges, and to broadly capture the extensive variation in PARylation levels. For instance, in the case of the FEN1 protein (PARylation stoichiometry = 0.031%), we were able to detect both the heavy and light species across all four mixing ratios, with the relative intensity of the heavy peak increasing as the mix ratio increased (Supplementary Table 1). Conversely, for the HNRNPUL1 protein (PARylation stoichiometry = 2.033%), which exhibited a PARylation occupancy 60 times higher than that of FEN1 according to our calculations, the heavy species was undetectable at the lowest mix ratio but became discernible at higher mix ratios as its relative intensity increased (Supplementary Table 1).

PARP1 emerged as one of the most heavily PARylated proteins with a PARylation stoichiometry of 32.9% under these conditions (Supplementary Table 2). Consistently, our measurements revealed the PARylation stoichiometry for several proteins known to undergo PARylation, including PARP1, YY1, CTCF, XRCC1, H2AFX, H2AFY, POLR1A, and POLR2A (Fig. 1G, and Supplementary Table 2)^22–25^.

To further assess the reproducibility of our findings, we plotted the median heavy-to-light peak area ratio for various proteins as a function of the mixing ratio. We found that the peak ratios and linearity for the same protein exhibited excellent agreement between the two replicate sets (Fig. 1H). Furthermore, we also performed independent biochemical experiments to validate our MS stoichiometry data. We generated lysates from H_2_O_2_-treated shPARG HCT116 cells, and isolated PARylated proteins using the boronate beads. The PARylated proteins were then eluted, and their abundances were compared to those in serial dilutions of the crude cell lysates. Our MS measurements indicated a PARylation stoichiometry of 0.63% for PCNA and 2.1% for EWSR1, which were consistent with the results obtained from the immunoblotting analysis (Fig. 1I).

### Networks of PARylated proteins with high PARylation stoichiometries

To further explore the potential roles of PARylation in DNA damage response, we analyzed the functional connections among the 235 identified PARylated proteins (Supplementary Fig. S1). Our analysis revealed that 71 proteins formed five networks, with functions associated with chromosome organization, regulation of gene expression, DNA repair, RNA metabolic process, and cell cycle, respectively (Fig. 2A).

**Fig. 2:**
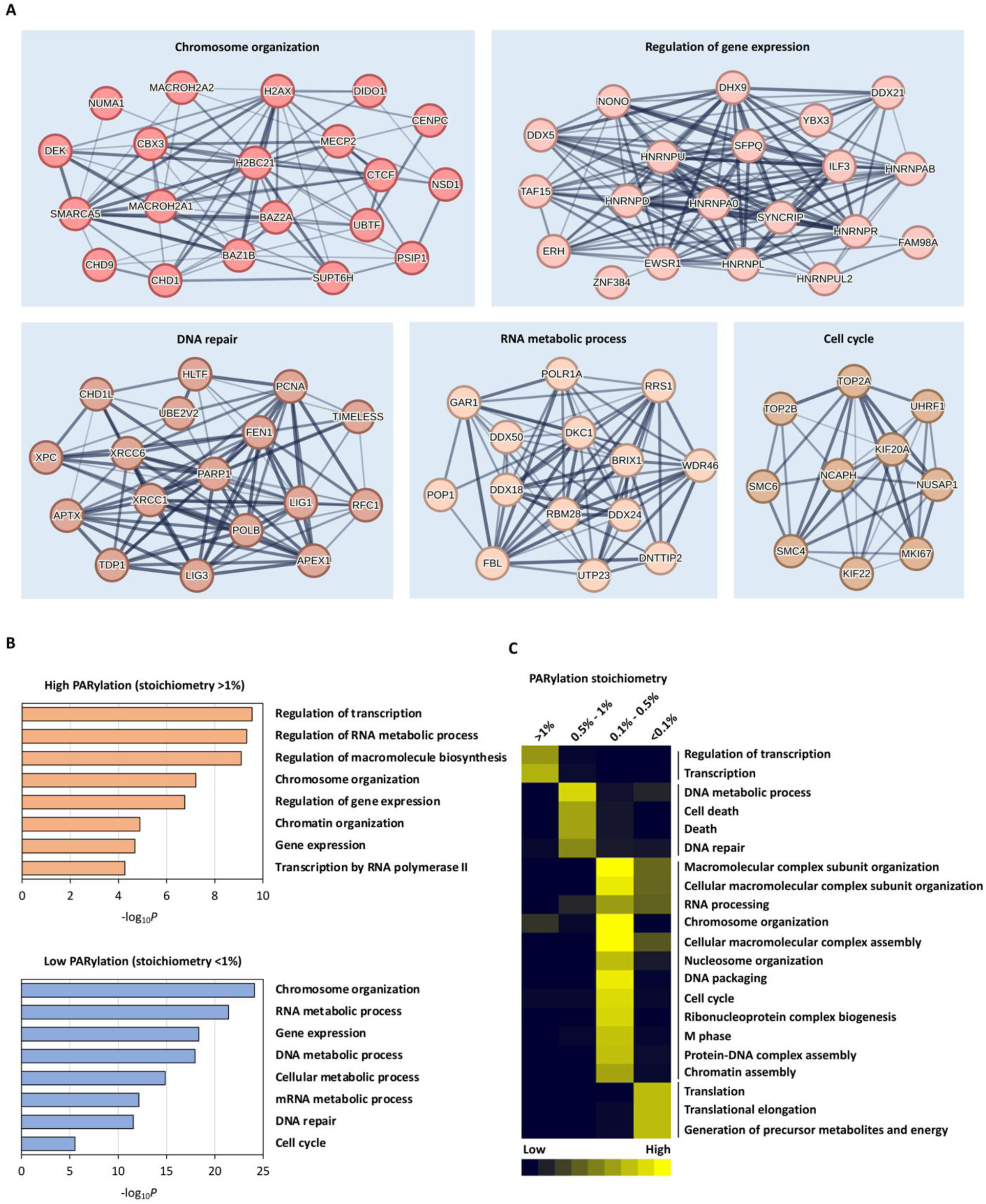
High-occupancy proteins are concentrated in transcription regulators, rather than DNA repair factors. **A** Network presentation of functional connections among the 235 PARylated proteins within the top five largest clusters identified through PARylation stoichiometry analysis. **B** Enriched biological processes in proteins with high (>1%) and low (<1%) PARylation stoichiometry. **C** Clustering of the enriched biological processes across different ranges of PARylation stoichiometry. The previously identified PARylated proteins (Table. S13, Zhang et al., *Nature Methods*)^13^ were used as a reference.

To gain further insights into the functional implications of PARylation stoichiometry, we conducted a GO analysis to identify enriched biological processes (BP) across different stoichiometry ranges (Fig. 2B, 2C). The PARylated proteins were categorized into four groups based on their PARylation stoichiometries: >1%, 0.5% - 1%, 0.1% - 0.5%, and <0.1%. It is worth noting that for the BP enrichment analysis, we employed the list of identified PARylated proteins as the background list, thus ensuring that the enriched processes were specifically associated with different stoichiometry values rather than being influenced by the entire human genome. Subsequently, we examined the enriched biological processes for each stoichiometry category and clustered them accordingly (Fig. 2C). Consistently, proteins with the highest occupancy (>1%) exhibited significant enrichment in GO terms related to transcription and regulation of transcription (Fig. 2C). Processes such as “DNA repair”, “DNA metabolic process” and “cell cycle” were enriched in proteins with PARylation stoichiometries between 0.5% and 1% (Fig. 2C). Our comprehensive analysis of PARylation stoichiometry thus indicates that high-occupancy proteins are predominantly concentrated in transcription regulators.

We further investigated the characteristics of proteins displaying the highest levels of PARylation occupancy (>1%) (Fig. 3A, and Supplementary Table 2). Notably, PARP1 emerged as a central hub protein, exhibiting the most extensive functional connections among the 91 highly PARylated proteins (Fig. 3A). This aligned with its well-known role as a key mediator of cellular PARylation. The list of proteins with high PARylation stoichiometries also included several well-established PARylated proteins, including YY1, TAF15, CHTOP, GAR1 and TIMELESS^13, 26–28^. Globally, the highly occupied (>1%) proteins were significantly enriched in transcription and regulation of transcription (Fig. 2C). Remarkably, within the top 91 proteins exhibiting the highest occupancy levels (>1%), as many as 57 proteins (62.64%) were identified as transcription regulators (Figure 3B).

**Fig. 3:**
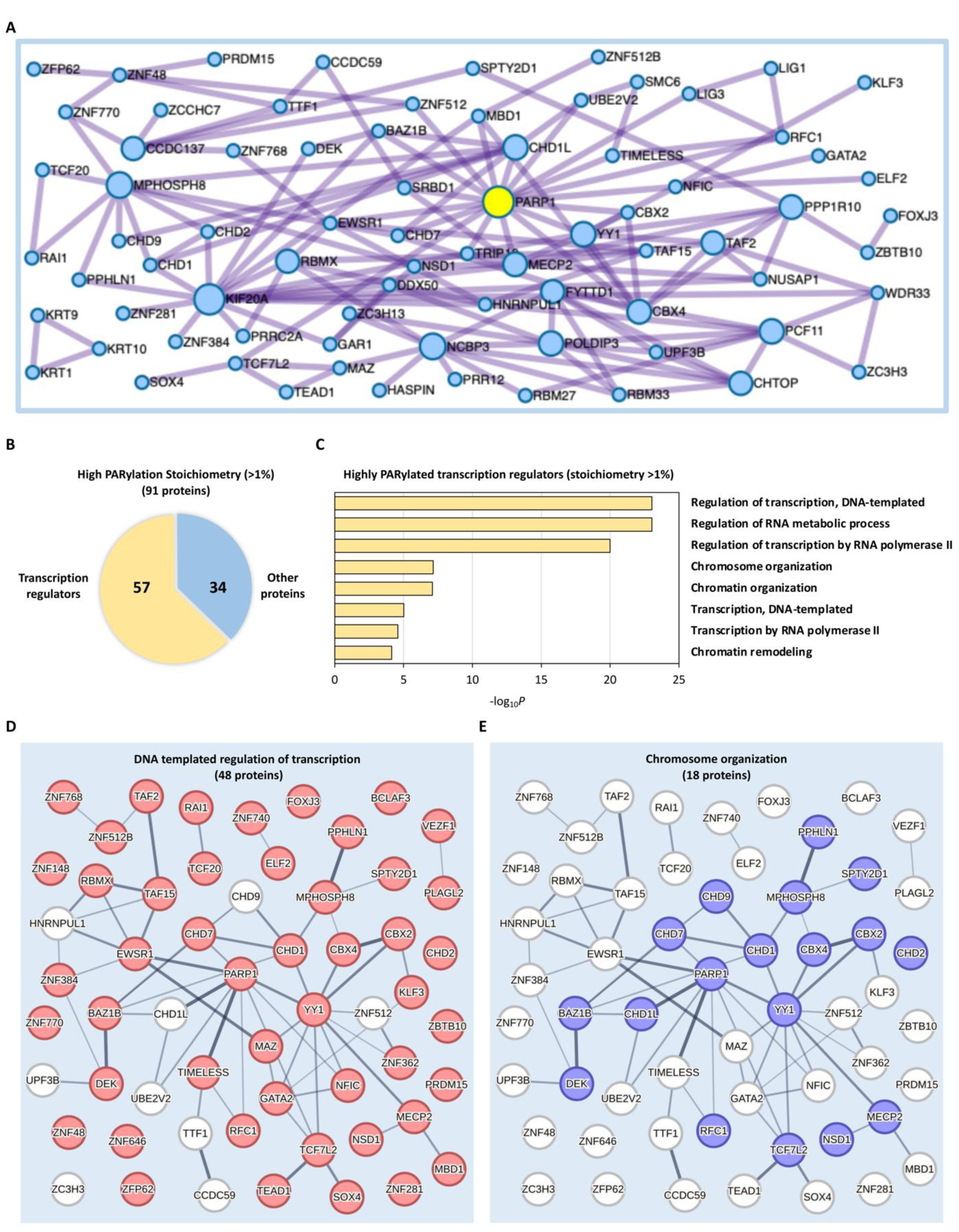
PARylation mainly regulates transcription during DNA damage response. **A** Protein-protein interaction enrichment analysis of the 91 highly PARylated proteins (stoichiometry >1%). **B** Proportion of transcription regulators among the top 91 highly PARylated proteins (stoichiometry >1%). **C** Enriched biological processes observed among the 57 transcription regulators within the top 91 highly PARylated proteins (stoichiometry >1%). **D, E** Network presentation of functional connections among the 57 highly PARylated transcription regulators within the two largest clusters identified through PARylation stoichiometry analysis. Proteins involved in DNA-templated regulation of transcription were marked in red (**D**), while proteins participate in chromosome organization were marked in blue (**E**).

The pronounced occupancy of PARylation in transcription regulators motivated us to delve deeper into their specific characteristics. To achieve this, we conducted a further GO analysis focusing on these highly PARylated transcription regulators (Fig. 3C). Furthermore, we found that these proteins formed two large networks, with functions predominantly associated with DNA-templated regulation of transcription (48 proteins) and chromosome organization (18 proteins), respectively (Fig. 3D, 3E). It has been well established that PARylation plays a central role in facilitating the recruitment of DNA repair factors to DNA lesions^29^. Our comprehensive PARylation stoichiometry analysis sheds new light on the significance of PARylation in controlling other biological processes in the nucleus, in particular transcriptional regulation and chromatin remodeling. This perspective expanded the functional repertoire of PARylation beyond the DNA repair factor recruitment.

### Transcription regulators have high PARylation stoichiometries

Our large-scale PARylation stoichiometry studies indicated that proteins involved in transcription regulation and chromatin remodeling have high PARylation stoichiometries. We further studied the 48 highly PARylated proteins (PARylation stoichiometries > 1%) that are involved in DNA-templated regulation of transcription. We found that more than 60% (29/48) of the proteins in this network were identified as transcription factors (Fig. 4A). Remarkably, protein domain analyses demonstrated that the majority (55.17%, 16/29) of these highly PARylated transcription factors are Cys2-His2 zinc finger (C2H2-ZF) proteins (Fig. 4B), which represent the largest group of putative human transcription factors (Fig. 4C, 4D). All these ZNF proteins are C2H2-type zinc finger transcription factors which target specific binding motifs on gene transcription starting sites (Fig. 4E, 4F). In this context, recent studies showed that proteins with C2H2-type zinc finger domains represent a major class of PAR-binding proteins^30, 31^.

**Fig. 4:**
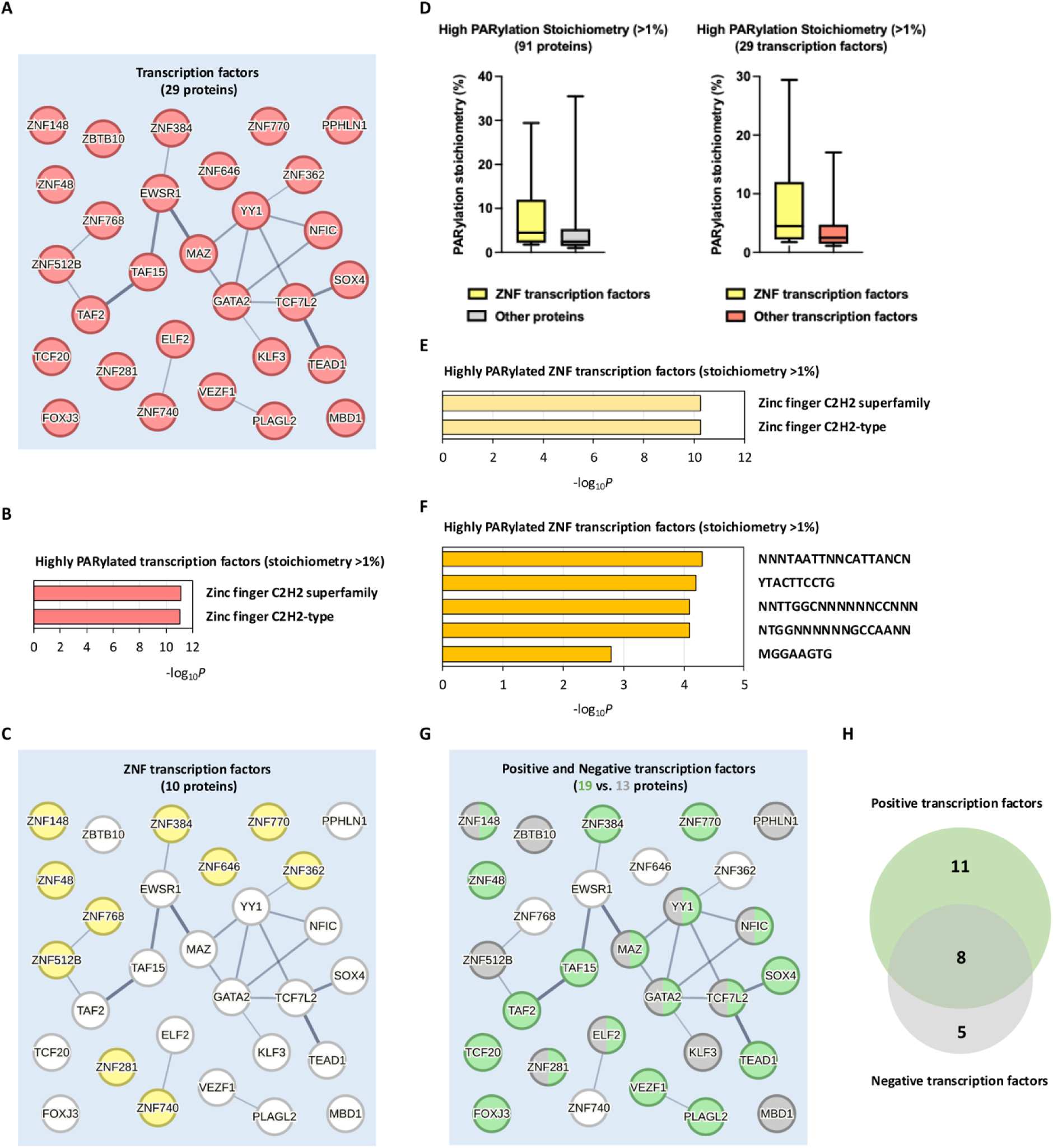
PARylation regulates transcription through modulating transcription factors. **A** Network presentation of functional connections among the 29 transcription factors within the 48 significantly PARylated transcription regulators (stoichiometry >1%) involved in DNA-templated regulation of transcription. **B** Protein domain analysis of the 29 highly PARylated transcription factors (stoichiometry >1%). **C** Network presentation of functional connections among a subset of highly PARylated ZNF transcription factors within the 29 highly PARylated transcription factors (stoichiometry >1%). **D** Distribution of PARylation stoichiometries among the ZNF transcription factors within the 91 highly PARylated proteins (stoichiometry >1%) (left), or within the 29 highly PARylated transcription factors (stoichiometry >1%) (right). The box represents the interquartile range, the middle line denotes the median, and the whiskers indicate the minimum and maximum values excluding outliers. **E** Protein domain analysis of the 10 highly PARylated ZNF transcription factors (stoichiometry >1%). **F** Transcription factor analysis of the 10 highly PARylated ZNF transcription factors (stoichiometry >1%). **G** Network presentation of functional connections among the positive (green) or negative (grey) transcription factors within the 29 highly PARylated transcription factors (stoichiometry >1%). **H** Biological processes analysis identified 19 positive transcription factors and 13 negative transcription factors among the 29 highly PARylated transcription factors (stoichiometry >1%). 8 multifunctional transcription factors were overlapped with both positive and negative properties.

These 29 highly PARylated transcription factors contained both positive (19 proteins) and negative (13 proteins) regulators of DNA-templated transcription (Fig. 4G, 4H). Among these proteins, we identified 8 multifunctional transcription factors, including YY1, GATA2, MAZ, NFIC, TCF7L2, ELF2, ZNF281, and ZNF148 (Fig. 4G, 4H). These 8 proteins can exhibit both positive and negative control over a substantial number of cellular genes by binding to their respective transcription start sites^32–47^.

For example, the transcription factor Yin Yang 1 (YY1) is a critical component of the RNA Pol II transcription machinery, and is a known PARylated protein (Fig. 4A)^22^. As a transcriptional repressor protein, PARylation of YY1 has been shown to trigger its dissociation from the PARP1 promoter binding site. This release of YY1 alleviates the repression of PARP1 transcription, which represents an important positive-feedforward mechanism to promote PARP1 expression and PARylation activity in response to cellular insults^33^. In our investigation, we observed that YY1 exhibited a remarkably high PARylation stoichiometry of 17.046%, under oxidative DNA damage conditions (Fig. 4A, and Supplementary Table 2). It is conceivable that PARylation stoichiometries could be a key factor that regulates the affinity of transcription factors towards their cognate DNA binding sites, thus mediating the expression of genes critical to DNA damage response.

In addition, TATA-box binding protein associated factor 15 (TAF15) is a transcription factor belongs to the FUS, EWS/EWSR1, TAF15 (FET) protein family^48, 49^. The FET family proteins play critical roles in transcriptional regulation, and their dysregulation through chromosomal translocations generates oncogenic fusion proteins that drive malignancies such as sarcomas and leukemias^50^. These fusion oncoproteins typically retain the N-terminal low-complexity domain (LCD) of FET proteins, which facilitates phase separation and transcriptional condensate formation, while incorporating the DNA-binding domain (DBD) of partner transcription factors, enabling aberrant gene regulation^50, 51^. For example, EWSR1-FLI1 is involved in the pathogenesis of Ewing’s sarcoma. It is formed as a result of a chromosomal translocation, which fuses the *EWSR1* gene’s N-terminal transcriptional activation domain with the C-terminal ETS-family DBD of *FLI1*^52, 53^. This chimeric protein acts as a potent transcriptional activator or repressor, depending on target gene context^52, 54^. Mechanistically, EWSR1-FLI1 undergoes phase separation at GGAA-rich loci, forming biomolecular condensates that recruit RNA polymerase II and coactivators, thereby amplifying transcriptional output^50^. Similarly, a TAF15 fusion protein is a result of a chromosomal rearrangement between the TAF15 gene and another transcription factor gene like *NR4A3* or *ZNF384*^55, 56^. These fusion proteins are found in sarcomas and some other cancers like extraskeletal myxoid chondrosarcoma or leukemia. They are considered strong transcription factors that can activate or repress genes^52–56^.

Furthermore, recent studies showed that all three FET proteins form pathological aggregates that are detected in various neurodegenerative diseases^57–61^. For example, abundant amyloid filaments of TAF15 were recently found in brain samples from frontotemporal dementia (FTD) patients^57^. The filament fold is formed from residues in the low-complexity domain (LCD) of TAF15, and is recurrently observed in TAF15 filaments residing in the motor cortex and brainstem of these patients. The formation of TAF15 amyloid filaments with a characteristic fold in FTLD establishes TAF15 proteinopathy in neurodegenerative disease^57^. In addition, EWSR1 is a prion-like protein, whose LC (low complexity) domains are well known to form stress-induced biomolecular condensates^62^. Importantly, *EWSR1* mutations have been identified in patients with familial ALS and FTD^63^. The ALS-associated mutant EWSR1 proteins exhibit an enhanced propensity for protein aggregation compared to WT EWSR1, suggesting that these mutations may precipitate accelerated aggregation within affected motor neurons^64^.

We previously found that TAF15 and EWSR1 were extensively PARylated with a total of 16 and 8 Asp-/Glu-PARylation sites, respectively^13^. Furthermore, PARylation of TAF15 and EWSR1 was highly sensitive to the treatment of PARP inhibitors (e.g., olaparib), suggesting that they are potential PARylation substrates of PARP1^13^. In this current study, we demonstrated that TAF15 is a highly PARylated protein with the PARylation stoichiometry of 7.278% under oxidative DNA damage conditions (Fig. 4A, and Supplementary Table 2). Furthermore, we found that EWSR1 (EWS) also exhibited high levels of PARylation (stoichiometry of 2.076%) under genotoxic stress conditions (Fig. 4A, and Supplementary Table 2). Importantly, it is increasingly appreciated that PARylation is a critical seeding mechanism that promotes the phase separation and aggregation of many intrinsically disordered proteins, including the FET proteins^65, 66^. It is conceivable that PARylation stoichiometry may serve as a critical determinant governing the spatiotemporal dynamics of these FET proteins, in the context of transcriptional regulation and biomolecular condensation.

### Chromatin remodelers have high PARylation stoichiometries

Our functional network analysis showed that a large number of highly PARylated transcription regulators are interconnected between DNA-templated regulation of transcription and chromosome organization (Fig. 3E and 5A). Notably, a majority (16 out of 18) of the highly PARylated chromatin remodelers directly participate in DNA-templated transcriptional regulation (Fig. 5B). This suggests that PARylation exerts its regulatory influence on multiple facets of the transcriptional processes through diverse mechanisms, including the modulation of chromatin remodeling.

**Fig. 5:**
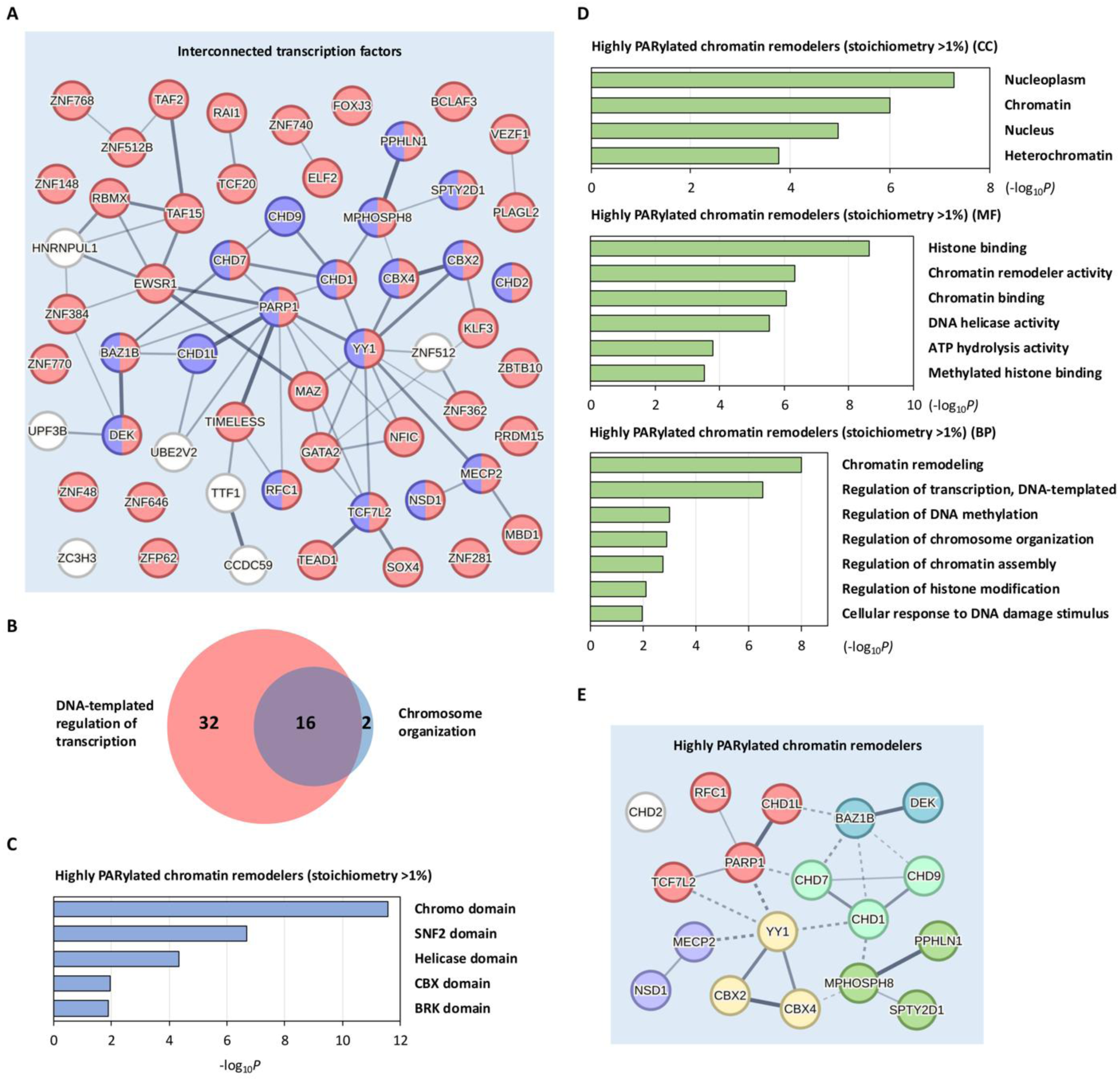
PARylation modulates transcription through chromatin remodeling. **A** Functional interconnection network of the highly PARylated transcription regulators (stoichiometry >1%) involved in DNA-templated regulation of transcription and chromosome organization. **B** Functional connection network analysis identified 16 highly PARylated transcription factors (stoichiometry >1%) involved in both DNA-templated regulation of transcription and chromosome organization. **C** Protein domain analysis of the 18 highly PARylated chromatin remodelers (stoichiometry >1%). **D** GO analysis of the 18 highly PARylated chromatin remodelers (stoichiometry >1%). CC, cellular component; MF, molecular function; BP, biological process. **E** Functional clustering networks of the 18 highly PARylated chromatin remodelers (stoichiometry >1%).

We conducted a protein domain analysis of the 18 highly PARylated transcription regulators involved in chromosome organization (Fig. 3E). Our analysis revealed a significant enrichment of Chromatin organization modifier domain (Chromo domain), Helicase domain, and SNF2 domain (ATPase domain) within these proteins (Fig. 5C). These findings align with the established functions of these proteins in chromatin remodeling^67–69^. Furthermore, GO analysis indicated that these highly PARylated substrates predominantly consisted of chromatin-associated proteins with chromatin remodeler activity and helicase activity involved in chromatin remodeling and DNA-templated regulation of transcription (Fig. 5D). Importantly, our functional network analysis indicated that these highly PARylated chromatin remodelers regulate various aspects of chromatin organization through diverse mechanisms, such as the regulation of chromatin accessibility, DNA methylation, and gene expression (Fig. 5E). Overall, these observations suggest that PARylation may regulate transcription through the modulation of chromatin remodeling processes.

Chromatin remodelers have recently been recognized as important signaling coordinators in DNA damage response^70^. Compelling evidence indicates that PARylation governs access and activities of certain chromatin remodelers, as exemplified by MECP2, CHD1L (ALC1) and DEK^67–69^. For instance, MECP2 is a chromosomal protein that promotes chromatin clustering. However, PARylation of MECP2 has a negative effect on high order chromatin structure, leading to chromatin decompaction^67^. We identified MECP2 as a highly PARylated protein (stoichiometry of 2.230%) in response to genotoxic stress treatments (Fig. 5E, and Supplementary Table 2).

PARylation is known to mediate the recruitment of different chromatin remodeling enzymes to the DNA damage site via their PAR-binding domains^71–73^. In this context, certain chromatin remodeling complexes can induce chromatin relaxation, enhancing DNA accessibility for transcriptional activation. One such example is the Chromodomain-Helicase-DNA-binding protein (CHD) 1-like (CHD1L, also known as ALC1), an SNF2-like ATPase that is recruited to DNA lesions by PAR through its macrodomain motif. This then triggers chromatin relaxation^68^, a process that is essential for effective transcriptional control^74^. Furthermore, PAR binding could also mediate the recruitment of repressive chromatin modifiers, where they bind to PAR and exert inhibitory effects on transcription. For example, the human oncoprotein DEK, a chromatin architectural protein, maintains a heterochromatic state in Drosophila through its interaction with PAR via its PBM domain^69^. Notably, we observed robust PARylation of both CHD1L (stoichiometry of 5.671%) and DEK (stoichiometry of 1.674%) in response to genotoxic stress treatments. These results suggest that direct PARylation of these proteins may also contribute to the functional regulation of their activities (Fig. 5E, and Supplementary Table 2). The presence of PAR moieties on these chromatin remodelers could also sterically disrupt protein-protein interactions (PPIs) and impede the formation of certain protein complexes during the chromatin remodeling process.

Collectively, our PARylation stoichiometry analysis provides compelling evidence that PARylation serves as a coordinator of multiple facets within the transcriptional regulation processes, employing diverse mechanisms such as the regulation of chromatin remodeling and the coregulation of transcription factors in response to genotoxic stress. These findings highlight the multifaceted role of PARylation in orchestrating the intricate network of regulatory events involved in maintaining genomic integrity and gene expression.

## Discussion

Stoichiometry (the fraction of a given protein that is modified at a given time) is a critically important parameter when assessing the functional importance of a protein posttranslational modification event. Wu et al.,^75^ previously developed an elegant strategy to globally measure protein phosphorylation stoichiometries. They utilized phosphatase treatment combined with LC-MS/MS analysis to determine proteome-wide phosphorylation stoichiometry^75^. Peptides resulting from whole cell lysates digestion were split in half. One fraction was treated with phosphatase, the other was not. The two fractions were chemically labeled with heavy and light tags respectively, mixed and analyzed by LC-MS/MS. The difference between the heavy and light species of a peptide containing known modification sites thus represents the phosphorylation stoichiometries. As another example, Weinert et al., presented a strategy for quantifying global acetylation stoichiometries^76^. Due to the very low stoichiometry of acetylation, direct comparison between enriched acetylated peptides and non-acetylated forms was not feasible. Instead, an in vitro chemical acetylation labeling method was employed, revealing that 86% of sites were acetylated at less than 1% occupancy.

However, characterization of the absolute stoichiometries of many other PTMs has been challenging^77^. This is particularly the case for PARylation. First, PARylation is a polymeric/heterogeneous modification that lacks a defined mass shift. As a result, direct detection of the PARylated species (and then comparing the ratio of the PARylated/unPARylated species to determine the PARylation stoichiometries) using MS-based methods is not feasible. Second, PARylation is of low abundance, and a robust enrichment strategy needs to be developed for its stoichiometric analyses. Finally, compared to other common PTMs, a rather unique feature of PARylation is that PARylation can occur on various amino acid acceptors, including Asp, Glu, Lys, Arg, Ser, Cys, His, and Tyr^29, 78^. It is important to note that these PAR-amino acid linkages display drastically different stability. The linkage between ribose and Ser (in Ser-ADP-ribosylation) is in an acetal moiety, and is chemically stable. In contrast, the bond between ADP-ribose and an Asp/Glu residue is within in a high energy hemiacetal ester moiety. As a result, Asp-/Glu-PARylation is uniquely susceptible to harsh experimental conditions, e.g., sample boiling ^13, 18, 79, 80^. In this context, some of the previous studies suggest that Ser-PARylation is the predominant form of PARylation in cells. However, harsh lysis conditions (sulfuric acid) were employed, which could lead to the artificial degradation of Asp-/Glu-PARylation sites and therefore cause bias^81^.

We previously developed an integrated strategy towards the site-specific analysis of the cellular Asp-/Glu-PARylated proteome^13^. These studies have led to the identification of thousands of unique, unambiguously assigned Asp/Glu-ADP-ribosylation sites^13, 82^. Here we leveraged this method and developed a large-scale method to measure the absolute stoichiometries of protein PARylation. As previously shown, because Asp-/Glu-PARylation is unstable, we used a mild lysis buffer (e.g., a neutral SDS buffer) to deactivate the PARP/PARG enzymes and to universally preserve the PAR chains. Importantly, this buffer system is compatible with the subsequent boronate enrichment procedures. In the current study, we further evaluated the boronate enrichment strategy. We found that it had extraordinary efficiency, which allowed a complete recovery of the PARylated proteins from crude cell lysates. This key feature of the boronate beads became the foundation of our PARylation stoichiometry approach (Fig. 1B). Specifically, we used the SILAC approach, and generated two samples, i.e., the light sample (with activated PARP1), and the heavy sample (with inhibited PARP1). The PARylated version of a protein in the light sample was recovered using boronate enrichment, and was digested. At the same time, the whole cell lysates of the heavy sample were digested to generate the peptides corresponding to the unPARylated version of this protein. By mixing the light peptides and heavy peptides, we were able to determine the absolute PARylation stoichiometries on the global scale. Because of the complete recovery of the PARylated proteins by the boronate affinity beads, this approach allows us to encompass all PARylation events occurring on diverse amino acid side chains.

In this study, we employed a Thermo LTQ-Velos Pro Orbitrap mass spectrometer for SILAC sample analysis, which offers a dynamic range of approximately 5,000^20^. However, it is important to note that proteins with very low or high PARylation stoichiometries could generate light or heavy peptides with extreme ratios that might fall outside the optimal dynamic range of the Orbitrap detector. To address this issue, we conducted a titration experiment, generating samples with varying heavy-to-light mix ratios (i.e., 1:50, 1:200, 1:800, and 1:2000). This approach ensured that SILAC peptide pairs with extreme light/heavy ratios under one SILAC mixture sample were recovered in subsequent samples within the titration series.

To investigate the influence of PARylation on protein function and regulation, we conducted an analysis of the enrichment of Gene Ontology categories across different stoichiometry ranges: high (>1%), medium (0.5% - 1%), low (0.1% - 0.5%), and very low (<0.1%). Clustering of biological processes based on PARylation stoichiometry values revealed several intriguing findings.

First, the majority of PARylation events occurred at very low levels. At the same time, PARylation stoichiometries also exhibited a wide dynamic range. This is consistent with that the synthesis of PAR polymers is highly regulated in a spatiotemporal manner. Upon binding to nicked DNA, PARP1 is rapidly activated, leading to the generation of many PARylated protein at the DNA lesions^18^. It has been estimated that the amount of PAR in a cell increases from ∼3,000 PAR molecules/cell under basal conditions, to >150,000 PAR molecules/cell under genotoxic conditions^83^.

Second, among all the measured proteins, PARP1 demonstrated one of the highest PARylation stoichiometric values (32.935%). This is consistent with the notion that a large fraction of the cellular PAR chains is attached to PARP1 itself^84^. Protein-linked PAR polymers (PAR chains) serve as a platform for the recruitment of DNA damage proteins (e.g., XRCC1, LIG3, and DNA Polβ) via their PAR-binding domains, which further promotes the formation of a large protein complex that mediates the repair of DNA breaks. Because PARP1 is a very abundant nuclear protein and it is PARylated with high stoichiometric values, it is expected that PARylated PARP1 plays a central role in acting as a scaffold to orchestrate the recruitment and coordination of DNA damage repair factors through its PAR chains.

Third, PARP1 activity influences several key DNA repair pathways, including base excision repair (BER), single-strand break repair (SSBR), nucleotide excision repair (NER), and nonhomologous end-joining (NHEJ)^85, 86^. In our dataset, we were also able to determine the PARylation stoichiometry of several proteins associated with these repair machinery (e.g., XRCC1, LIG3, PCNA, XPC, and XRCC6) (Fig. 2A). These results suggest that besides PAR-binding, direct PARylation of these DNA repair factors could also facilitate DNA repair processes. However, we also observed that many of these DNA damage repair factors are associated with low PARylation stoichiometries (Supplementary Table 2). These results suggest that there could be a complex interplay between PAR-binding vs. direct PARylation for these DNA repair proteins (Fig. 2A). Therefore, whether PARylation regulates the activity of these DNA repair proteins via protein-protein interactions vs. covalent PARylation will be examined in future studies.

Fourth, many proteins with the highest stoichiometries were associated with the biological processes of “Regulation of Transcription”. This observation suggests that a significant fraction of these transcription regulator molecules is in a pool that is accessible to PARP1, which become PARylated under genotoxic conditions. PARP1 is known to act as a positive cofactor in transcription. At the same time, its activation can also repress RNA polymerase II-dependent transcription^87, 88^. Previous studies have demonstrated that PARP1-mediated transcriptional silencing involves the PARylation of the TATA-binding protein^88^. Notably, this modification effectively prevents the formation of active transcription complexes (e.g., for the transcription factor YY1^33^). However, once transcription complexes are assembled, the initiation of PARylation no longer affects the DNA binding of transcription factors. Consequently, if transcription factors are already bound to DNA, they become inaccessible to PARylation^89^. Hence, PARylation inhibits the binding of transcription factors to DNA, while DNA binding prevents their modification. It is plausible to propose that PARylation serves as a molecular switch to block the DNA binding of transcription factors, thereby preventing the expression of damaged genes. In our current study, we identified eight highly PARylated multifunctional transcription factors, exhibiting significant PARylation stoichiometries under the oxidative DNA damage conditions (Fig. 4H,4I, and Supplementary Table 2).

Finally, proteins with high PARylation stoichiometries were also enriched with chromatin remodelers. It is plausible that PARylation reconfigures chromatin structure, resulting in altered chromatin accessibility and the exposure of DNA, which provides accessible binding sites for transcription factors and DNA repair proteins. This highlights the significance of PARylation-mediated transcriptional regulation by coordinating transcription factors and chromatin remodelers during DNA damage response.

PARylation has garnered considerable attention, particularly due to tremendous success of PARP inhibitors in the clinic^90^. Previously, it was summarized that PARylation regulates the first wave of the DNA damage response^29^. Understanding the stoichiometry of PARylation represents an important parameter to evaluate the impact of PARylation on protein function. We highlighted several protein classes with high PARylation stoichiometries. Among the highly PARylated transcription regulators are a class of poorly studied ZNF proteins. Our analysis revealed substantial PARylation occupancy in these ZNF proteins. This finding implies that the direct PARylation of these proteins may play a crucial role in regulating their functional properties. Interestingly, a recent study showed that many of these ZNF proteins also bind to PAR polymers^31^. Whether the PARylation of these transcription factors regulates their function positively or negatively and how specific types of transcription factors are involved in DNA damage response coupled transcriptional regulation remain to be elucidated.

In addition, we demonstrated that TAF15 and EWSR1, two members of the FET proteins, were extensively PARylated with high stoichiometric values. The FET proteins are DNA/RNA binding proteins that are implicated in DNA damage response and transcription regulation. When fused with another gene (FLI1, ATF1, CREM, RBFOX2, ERG, TCF7L2, TFEB and ZNF384), they form abnormal chimeric transcription factors (e.g., EWSR1-FLI1, EWSR1-ATF1 and TAF15-ZNF384) that drive the pathogenesis of a variety of soft tissue tumors, leukemias and mesotheliomas^55, 91, 92^. Thes results point to the intriguing possibility of the regulatory importance of PARylation in modulating DNA damage response through transcriptional regulation.

Recent studies showed that the FET proteins form pathological aggregates that drive the pathogenesis of various neurodegenerative diseases^57–61^. Indeed, mutations of the FET proteins are identified in patients with familial ALS and FTD, and these mutations greatly accelerate the formation of fibrils^59, 93, 94^. Abundant amyloid filaments of TAF15 were recently found in brain samples from FTD patients^57^. It has been shown that the low complexity domain in these FET proteins function as prion proteins that drive the formation of biomolecular condensates. Furthermore, our previous PARylation proteomic studies identified a total of 16 and 8 PARylation sites on TAF15 and EWSR1, respectively. Many of the sites on these FET proteins were exquisitely sensitive to PARP inhibitor treatment, indicating that they are novel PARP1 downstream targets^13^. The RRM (RNA Recognition Motif) in these proteins are known to bind to PAR polymers^95^. Because these proteins are PARylated with high PARylation stoichiometries, the high local concentrations of PAR polymers in combination with the PAR-binding domains in these proteins could function as a particularly important seeding mechanism (i.e., via multivalency) to promote their aberrant biomolecular condensation. This important aspect of PAR biology in the context of neurodegenerative diseases warrants future studies.

Our study offers a large-scale method to measure absolute PARylation stoichiometries. At the same time, it is also important to acknowledge its limitations. Because we used boronate beads to pull down the PARylated proteins, this method allows for the determination of PARylation stoichiometries at the protein level, but not specific PARylation sites. Future methodologies are needed to assess stoichiometries at the level of individual PARylated residues. Here, we describe a novel large-scale approach for the measurement of absolute PARylation stoichiometries. Because PARylation is a labile, heterogenous and low-abundance modification, we performed extensive optimization of our approach, including cell lysis conditions, boronate enrichment efficiency and carefully designed titration experiments. In total, we were able to measure the absolute PARylation stoichiometries of 235 proteins under genotoxic conditions. Our analysis revealed that protein PARylation stoichiometry varies significantly among PARylated proteins, with pronounced dependence on specific biological processes. Functional analyses of this dataset revealed a remarkable breadth of regulatory events associated with the PARylated proteins, including DNA damage repair but also RNA metabolism, transcriptional regulation and chromatin remodeling. We unveiled that proteins with high PARylation stoichiometries are involved in coordinating various aspects of transcriptional regulation and chromatin remodeling during DNA damage response. The quantitative proteomics approach and the associated datasets described herein represent a valuable resource for system-level investigations into the functions of protein PARylation dynamics.

## Materials and Methods

### Cell culture and reagents

HCT116 cells with PARG knockdown were generated following previously described methods^13^. The HCT116 shPARG cells were cultured in SILAC medium. SILAC Dulbecco’s Minimal Essential Medium (DMEM) were supplemented with 10% dialyzed FBS and 2 µg/ml puromycin. To prepare the SILAC media, 84 µg/ml light Arginine was added to the medium, and 152 µg/ml light or heavy Lysine was added to prepare the light or heavy DMEM media. Anti-poly-(ADP-ribose) antibody was obtained from Tulip Biolabs (#1020). PCNA (#2586), PARP1 (#9542) and GAPDH (#5174) antibodies were purchased from Cell Signaling Technology. EWSR1 antibody (#A300-417A) was purchased from Bethyl Laboratories. *M-aminophenyl-boronic* acid–agarose (A8312) was obtained from Sigma. Unless specified, all other chemicals and reagents were also obtained from Sigma.

### Boronate affinity beads (BAB) pulldown

The purification of PARylated proteins for immunoblotting analysis was conducted following the previously described method^13^. In brief, H_2_O_2_ stimulated cells were lysed using SDS lysis buffer (1% SDS, 10 mM HEPES, pH 7.0, 2 mM MgCl_2_, 250 U universal nuclease). The cell lysates were then subject to BAB enrichment at room temperature for one hour. Subsequently, fresh BAB was added to the mixture and rotated for an additional hour. Crude lysates, as well as the supernatants from the first and second rounds of adsorption, were collected and subjected to immunoblotting analysis to demonstrate the quantitative adsorption of PARylated proteins. Immunoblotting was also used to validate our stoichiometry measurement. The PAR modified proteins were eluted from the BAB using 3 M NH_4_Ac, pH 5.0, and the eluted protein solution was diluted with SDS lysis buffer to the same volume as the starting cell lysates. Meanwhile, a fraction of the cell lysates was diluted to 1/25 ∼ 1/200 of the original concentration using SDS lysis buffer. Subsequently, equal volumes of the serially diluted cell lysates (input) and the pull-down eluates were analyzed by immunoblotting.

### Immunoblotting analysis

The samples were subjected to electrophoresis using the standard SDS-PAGE method. Subsequently, the proteins were transferred onto a nitrocellulose membrane (Whatman). The membranes were then blocked with a TBST buffer (25 mM Tris-HCl, pH 7.5, 150 mM NaCl, 3% BSA) and incubated overnight at 4 °C with primary antibodies. Following this, peroxidase-conjugated secondary antibodies were applied for 1 hour at room temperature. The blots were developed using enhanced chemiluminescence, exposed on autoradiograph films, and processed using standard methods.

### Sample preparation for stoichiometry measurement by MS

Light Lys and heavy Lys-labeled shPARG HCT116 cells were treated with H_2_O_2_ (2 mM, 5 min) or Olaparib (1 µM, 40 min) respectively. Both light and heavy cells were lysed using SDS lysis buffer. The protein was quantified using a BCA assay kit, followed by reduction, alkylation, and chloroform/methanol precipitation. The heavy lysates protein was resuspended in urea (pH 7.5) to a final concentration of 250 µg/ml and digested overnight with Lys-C at a 1:100 enzyme-to-protein ratio. Simultaneously, approximately 65 mg of light lysates protein was enriched for PARylated proteins using BAB according to the previously described method section. The protein-bound BAB beads were extensively washed using SDS wash buffer (1% SDS, 100 mM HEPES, pH 8.5, 150 mM NaCl), followed by 100 mM HEPES (pH 8.5) and 150 mM NaCl to remove any residual detergent. Finally, the beads were resuspended in 200 mM HEPES pH 8.5, and the proteins were on-bead digested overnight with Lys-C. After digestion, aliquots of light and heavy peptides were mixed to different ratios of crude lysates protein from which the peptides are derived:

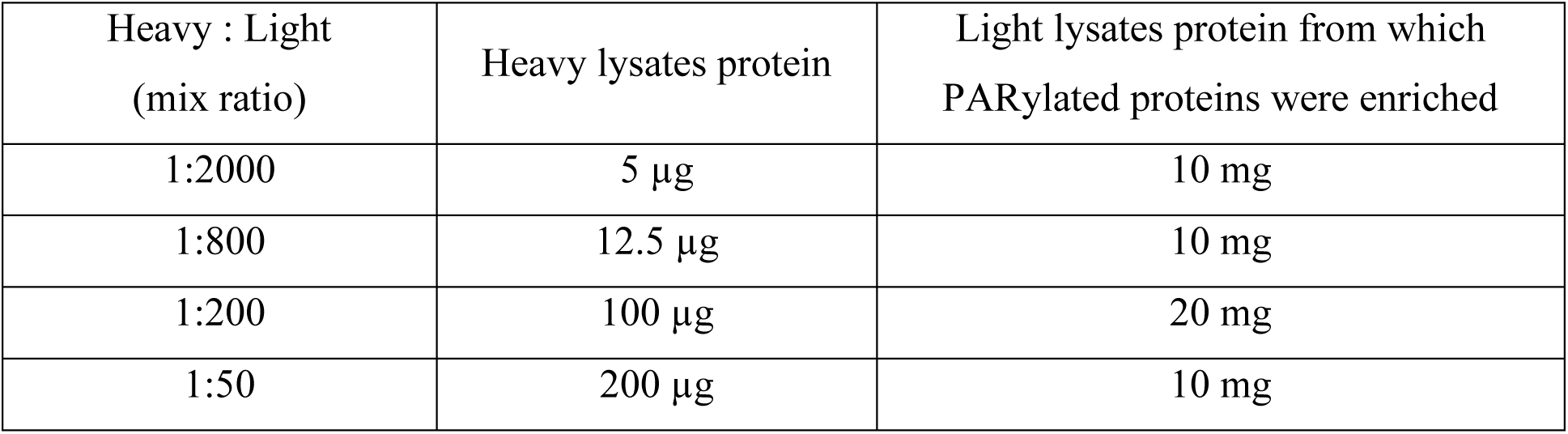

| Heavy : Light<br>(mix ratio) | Heavy lysates protein | Light lysates protein from which<br>PARylated proteins were enriched |
| --- | --- | --- |
| 1:2000 | 5 μg | 10 mg |
| 1:800 | 12.5 μg | 10 mg |
| 1:200 | 100 μg | 20 mg |
| 1:50 | 200 μg | 10 mg |

Depending on the peptide abundance, the mixtures were either directly analyzed using LC-MS/MS or further fractionated using Basic pH RP-HPLC prior to LC-MS/MS analysis. For the samples with ratios of 1:2000 and 1:800, the peptide mixtures were desalted and directly subjected to LC-MS/MS analysis. However, for the samples with ratios of 1:200 and 1:50, the peptide mixtures were desalted and further fractionated using basic pH RP-HPLC (ZORBAX 300Extend-C18, 3.5µ).

A gradient elution was performed over 90 minutes, starting from 95% buffer A (10 mM ammonium formate, pH 10.0) to 90% buffer B (10 mM ammonium formate, 90% ACN, pH 10.0). Fractions were collected every 1.5 min from 5 min to 80 min, and subsequently concatenated into 17 final fractions based on the scheme presented by Wang et al.,^96^. These fractions were then desalted and subjected to LC-MS/MS analysis.

To quantitatively evaluate the non-specific binding of the non-PARylated form of proteins to the BAB, we employed SILAC-labeled HCT116 shPARG cells. The cells labeled with heavy Lysine were treated with Olaparib (1 µM, 40 min), while the cells labeled with light Lysine were treated with H_2_O_2_ (2 mM, 5 min). Subsequently, 5 mg of heavy cell lysates and 5 mg of light cell lysates were combined and subjected to boronate beads enrichment, followed by on-beads digestion. The resulting mixture of peptides was desalted and analyzed by MS. In this case, the heavy peak represents the non-specific binder, and the ratio of the heavy peak to the light peak of peptides derived from a PARylated protein was utilized to quantify the extent of non-specific binding.

### Mass spectrometry analysis and data processing

Peptide quantification was carried out following the established protocol described previously^13^. The peptide list was filtered based on the following criteria: (1) the signal to noise ratio of either heavy or light peak must be higher than 5; (2) peptide must be from known PARylated proteins (i.e., present in Table. S13, Zhang et al.,^13^); and (3) peptides containing known PARylation sites must be excluded. Median of the ratios of heavy peak area to light peak area was subsequently calculated for peptides derived from the same protein.

PARylation Stoichiometry at a mix ratio = (mix ratio) / (heavy/light MS peak area ratio median) The median of stoichiometries obtained from all mix ratios was taken as the final PARylation stoichiometry of the protein.

### GO pathways enrichment-based clustering of proteins

The proteins were categorized into four groups based on their PARylation stoichiometry: <0.1%, 0.1% - 0.5%, 0.5% - 1%, and >1%. For each group, enrichment analysis of the GO^97^ Biological Processes was conducted separately using DAVID software^98^, with the entire list of PARylated proteins serving as the background (Table. S13, Zhang et al.,^13^). The resulting *P*-values for each enriched (*P* < 0.05) were transformed into x = −log_10_(*P*), and these x-values for all four groups were subjected to hierarchical clustering (Euclidean distance, Single linkage) using Cluster 3.0 software^99^. The clustering results were then visualized using Java TreeView software^100^.

## Acknowledgments

This work was supported by the grant from the NIH (5R35GM134883). We thank Dr. Shan Zha and members of the Yu lab for helpful discussions.

---

**Figure S1.**
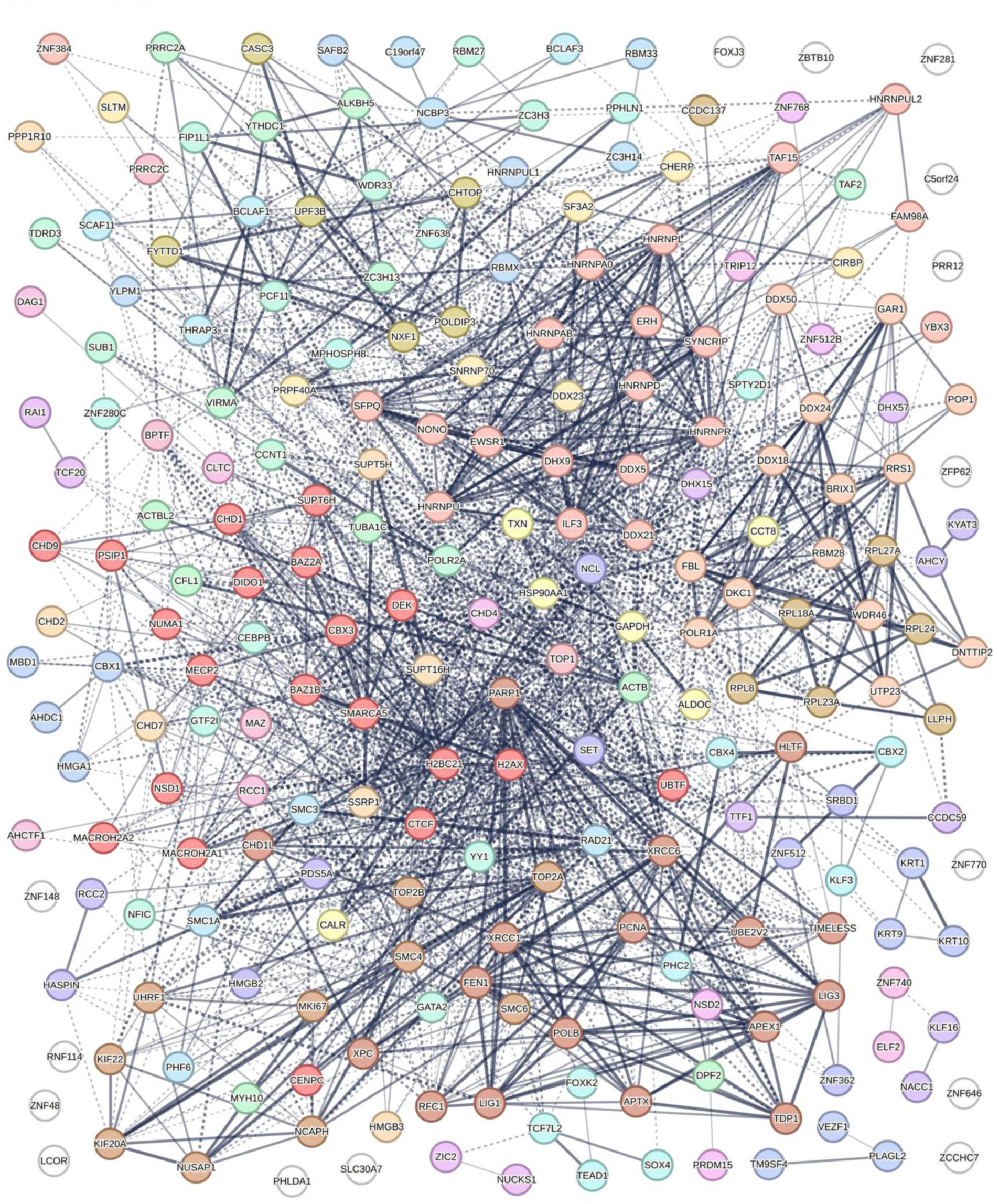
Functional connections among the PARylated proteins. The network of functional connections among the 235 PARylated proteins identified in PARylation stoichiometry analysis was determined by STRING analysis (STRING Version 11.5).

